# Postnatal maturation of serotonergic modulation of spinal RORβ interneurons in the medial deep dorsal horn

**DOI:** 10.1101/2024.11.15.623295

**Authors:** Christopher W. West, D. Leonardo Garcia-Ramirez, Kimberly J. Dougherty

## Abstract

Proprioceptive input is essential for coordinated locomotion and this input must be properly gated to ensure smooth and effective movement. Presynaptic inhibition mediated by GABAergic interneurons provides regulation of sensory afferent feedback. Serotonin not only promotes locomotion, but also modulates feedback from sensory afferents, both directly and indirectly, potentially by acting on the GABAergic interneurons that mediate presynaptic inhibition. Developmental disruptions in presynaptic inhibition can produce deficits in sensorimotor processing. Importantly, both presynaptic inhibition of proprioceptive afferents and serotonergic innervation of the spinal cord become mature and functional after the first postnatal week. However, little is known about the serotonergic receptors involved in the modulation of interneurons mediating presynaptic inhibition and when developmentally their actions mature. Here, we used whole-cell patch clamp recordings in lumbar spinal slices from neonatal and juvenile mice to assess the intrinsic properties and serotonergic modulation of deep dorsal horn GABAergic RORβ interneurons previously shown to mediate presynaptic inhibition of proprioceptive afferents. RORβ interneurons from juvenile cords displayed more mature membrane properties. Further, serotonin increased the excitability of RORβ interneurons via actions at 5-HT_2A_, 5-HT_2B/2C_, and 5-HT_7_ receptors in juvenile but not early neonatal spinal cords. Our findings indicate that deep dorsal horn RORβ interneurons undergo postnatal maturation in both their intrinsic excitability and ability to respond to serotonin, concurrent with the maturation of serotonergic innervation of the dorsal horn. This information can prompt future targeted studies testing relationships between impairments of serotonergic development, proprioceptive processing disorders, and presynaptic inhibition mediated by RORβ interneurons.

## Introduction

For humans and many other mammals, the development of efficient and coordinated locomotion involving weight-bearing stepping and postural control matures postnatally (1–3). In mice and rats, weight-bearing locomotion does not develop until the second postnatal week (4–7). However, much of our knowledge about the organization of locomotor circuits, neuronal types involved, and molecular fingerprints of those neurons comes from work in early neonatal rodents (8–10). The spinal neuronal circuits which generate locomotion are present and functional at birth (4, 5, 8, 11). At least at the level of the central pattern generator, the circuit organization seems relatively maintained into adulthood, as demonstrated by the functional consequences of genetic removal or silencing of specific neuronal types (9, 10, 12, 13). However, there are changes in the intrinsic and synaptic operations within the circuit over the first postnatal weeks (14, 15).

Both descending commands and afferent feedback are essential for coordinated weight-bearing locomotion (16–22). Afferent input conveys necessary sensory feedback from the environment to the spinal cord constantly during locomotion. This proprioceptive and mechanoreceptive input barrages the cord and must be properly controlled to ensure smooth locomotion (17, 22). Within the spinal cord, afferent transmission and reflexes are regulated by inhibition, and one of the most prominent forms of regulation is presynaptic inhibition (23–26). Presynaptic inhibition is predominantly produced by a trisynaptic pathway where sensory afferents activate excitatory interneurons which then activate GABAergic interneurons that release GABA back onto the afferents, leading to a shunting of afferent transmission (19, 23, 25). Although sensorimotor reflexes are present at birth (27, 28), GABAergic presynaptic inhibition of proprioceptive afferents becomes functional after the first postnatal week (29). GABAergic terminals on proprioceptive afferents are detected at P3 and increase over the first postnatal weeks (20, 28). Loss of presynaptic inhibition leads to perturbed movement (17) and loss of presynaptic inhibition of proprioceptors by medial deep dorsal GABAergic interneurons identified by RORβ expression specifically, results in an abnormal gait characterized by hyperflexion (21).

Another essential component for the maturation of locomotion is serotonin, released exclusively by descending supraspinal projection neurons (5, 30–33). Serotonergic innervation of the lumbar spinal cord matures postnatally for several weeks (34, 35), coincident with the maturation of the GABAergic presynaptic inhibition of sensory afferents (29, 35), as well as the development of weight bearing stepping in mice (4–7). Importantly, impairments in serotonergic modulation during development are associated with deficits in sensory gating (36), and the development of autism (37). Impaired presynaptic inhibition plays a role in sensory processing difficulties related to autism spectrum disorder (38, 39). Serotonin modulates presynaptic inhibition (40–43), although it is unknown exactly where in the trisynaptic loop this occurs aside from at the level of the afferents (43–47).

Thus, the goal of this study is to determine the early postnatal maturation of the intrinsic properties and serotonergic control of medial deep dorsal horn GABAergic interneurons mediating presynaptic inhibition of proprioceptive afferents, genetically identified by expression of RORβ (21). We hypothesized that serotonin has predominantly excitatory effects on RORβ interneurons, but these effects are strengthened postnatally due to differences in the intrinsic properties and expression of activatable serotonin receptors. Our data demonstrate that RORβ interneurons from neonatal and juvenile mice differ in their intrinsic electrophysiological properties, and in the strength of response to serotonin. We find juvenile RORβ interneurons have greater excitability overall, as well as excitatory responses to serotonin and to the activation of select serotonin receptors. Our results indicate that both intrinsic excitability and serotonergic modulation of deep dorsal horn RORβ interneurons increase with postnatal maturation. These findings can inform future investigations targeting serotonergic mechanisms underlying sensorimotor developmental disorders and possibly spinal cord injury.

## Materials and Methods

### Mouse lines and experimental groups

Experiments were performed using male and female *RORβ::cre*; *Rosa26-flox-stop-flox-tdTomato* (Ai9) mice (Jax mice #023526 and #007909, respectively) (48, 49). All experimental procedures followed National Institutes of Health guidelines and were approved by the Institutional Animal Care and Use Committee at Drexel University. Experiments were performed using two age groups: (1) neonatal (P1-5; N=23) and (2) juvenile (P14-21; N=22).

### Spinal Cord Preparations

For electrophysiology experiments, juvenile mice were first anesthetized, with ketamine (150 mg/kg) and xylazine (15 mg/kg). Neonatal and juvenile mice were decapitated, eviscerated, and cords were dissected. Dissection and slicing solutions were ice cold and contained: 111mM NaCl, 3mM KCl, 11mM Glucose, 25 mM NaHCO_3,_ 3.7mM MgSO_4_, 1.1mM KH_2_PO_4_, 0.25mM CaCl_2_. For mice >P7, 222mM glycerol was substituted for the NaCl to promote viability of the older cord (50, 51). Lumbar spinal cord spanning from L1-L4 was removed, embedded with 3% agar, then sectioned transversely (300µm) using a Leica Microsystems vibrating microtome. Spinal slices were transferred to oxygenated, room temperature, artificial cerebrospinal fluid (ACSF), containing: 111 mM NaCl, 3mM KCl, 11mM Glucose, 25 mM NaHCO_3,_ 1.3 mM MgSO_4_, 1.1mM KH_2_PO_4_, 2.5mM CaCl_2_. All solutions were continuously bubbled with 95% O_2_ and 5% CO_2_.

### Whole Cell Patch Clamp Recordings

Patch clamp recordings were performed in room temperature ACSF. RORβ interneurons fluorescently labeled by expression of TdTomato were visualized using a 63X objective lens on an Olympus BX51WI scope using LED illumination provided by a Lumen Dynamics X-Cite Fluorescence Illumination System. RORβ interneurons were selected for recording based on their anatomical localization in medial deep dorsal horn, which was confirmed by visualizing the placement of the recording electrode with a 5x objective after the conclusion of recording. Electrodes were pulled using a Sutter Instruments multistage puller to have tip resistances of 5-8 MΩ. Electrodes were filled with an intracellular solution containing: 128mM K-gluconate, 10mM HEPES, 0.0001mM CaCl2, 1mM glucose, 4mM NaCl, 5mM ATP, and 0.3mM GTP. Electrophysiology data was recorded using Molecular Devices Multiclamp 700B amplifier and Clampex software pClamp9. Recordings were digitized at a sample rate of 20 kHz and filtered at 4 or 6kHz.

### Agonist and antagonist application

All drugs utilized in these experiments were purchased from Sigma Millipore. Serotonin and receptor agonists and antagonists were made as stock solutions (1-10mM) and stored at - 20°C until needed for experiments. SB206553 was dissolved in DMSO and diluted to a final DMSO concentration of 0.02%. All drugs were bath applied for 10-15 minutes prior to recording intrinsic properties. Serotonin was applied in ACSF at a concentration of 10 µM. To determine the presence and role of select 5-HT receptors on RORβ interneurons, intrinsic properties were measured at baseline (ACSF alone), 10-15 minutes after bath application of a 5-HT receptor antagonist (SB206553, ketanserin, or WAY-100635), and then 10-15 minutes after bath application of 5-HT receptor agonists (DOI or 8-OH-DPAT). Serotonin receptor antagonists/agonists were applied in ACSF at the following concentrations: SB206553 (1 µM), ketanserin (1 µM), WAY-100635 (10 µM), DOI (10 µM), and 8-OH-DPAT(10 µM).

### Data Analysis

Analysis was performed using Clampfit software, and GraphPad Prism was used to perform statistical tests, analyses, and data visualization. All data was tested for normality in the distribution using Shapiro-Wilks test. If data passed normality tests, t-tests (paired or unpaired) were performed, depending on the experiment. Unpaired t-tests were used for data comparing neonate to juvenile intrinsic properties, while paired t-tests were used for before and after comparisons for pharmacology experiments. When data did not pass normality test, Mann-Whitney U test and Wilcoxon matched pairs signed rank test (before and after drug conditions) were used. Statistical significance was set to *p*<0.05. Results are reported as mean +/− standard deviation.

## Results

### Intrinsic properties of neonatal and juvenile RORβ interneurons

We first determined the intrinsic electrophysiological properties of RORβ interneurons in neonatal (P1-5) and juvenile (P14-21) mice. We performed whole cell patch clamp recordings of visually identified RORβ interneurons in the medial deep dorsal horn (lamina IV-VI) from lumbar sections of spinal cords from *RORβ::cre*; *Rosa26-lsl-tdTomato* mice (Fig. 1). Four firing patterns were observed in response to depolarizing current steps: tonic, initial burst, delay and single spike. In neonatal mice, 43% of RORβ interneurons recorded displayed tonic firing, 13% exhibited initial burst firing, 25% displayed delayed firing, and 19% fired single spikes (Fig. 2A). In contrast to neonatal RORβ interneurons, 61% of juvenile RORβ interneurons fired tonically, 22% were initial burst firing, and 17% fired with a delay (Fig. 2A). None of the juvenile RORβ interneurons recorded fired with a single spike. These data show that as RORβ interneurons mature, there is an increase in tonic firing and a decrease in the single spiking neurons.

**Figure 1.**
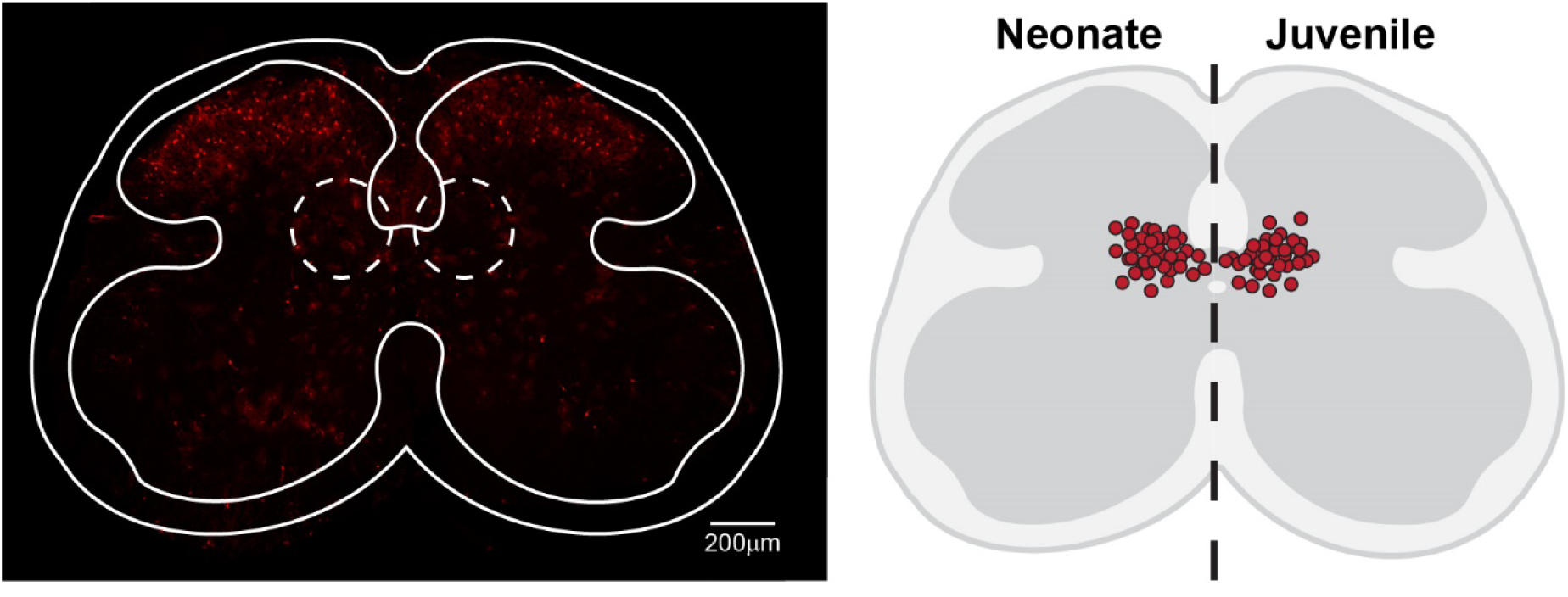
Spinal slice preparation used for whole cell patch clamp experiments. Left: Representative spinal slice showing the distribution RORβ interneurons (in red) in the medial deep dorsal horn (dotted circles). Right: Diagram displaying the locations of RORβ interneurons recorded from in neonatal (n=33) and juvenile mice (n=32).

**Figure 2.**
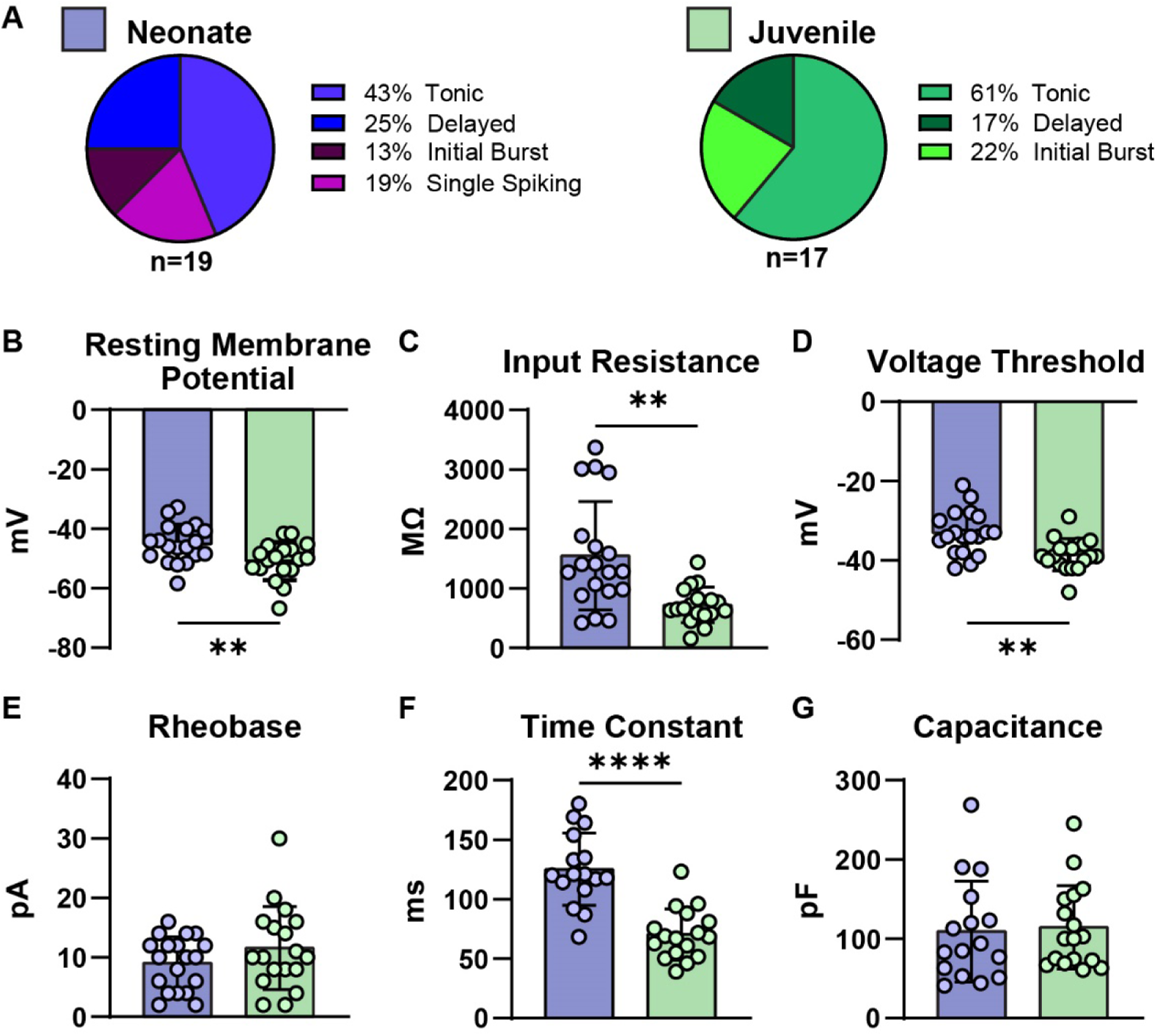
Intrinsic electrophysiology properties of RORβ Interneurons differ in postnatal developmental stages. (A) The observed firing patterns of RORβ interneurons in neonate and juvenile mice. (B-G) Comparison of membrane properties between RORβ interneurons from neonatal (blue) and juvenile (green) mice: (B) Resting membrane potential, (C) input resistance, (D) voltage threshold for an action potential, (E) rheobase, time constant (F), and capacitance (G). *=p<0.05, **=p<0.01, ****=p<0.0001; *t-test* or Mann-Whitney U-Test.

In addition to a shift towards tonic firing, commonly observed in mature inhibitory neurons in the dorsal horn (52). We found differences in the excitability of RORβ interneurons from neonatal and juvenile mice. There was a significant hyperpolarization in the resting membrane potential (neonate: −45.1±6.4 mV; juvenile: −50.8±6.3 mV, p= 0.009; Table 1; Fig. 2B), decrease in input resistance (neonate: 1551±911 MΩ; juvenile: 718±298 MΩ; p=0.001; Table 1; Fig. 2E), hyperpolarization of voltage threshold for an action potential (neonate: −33.1±5.5 mV; juvenile:-38.6±4.0 mV, p= 0.002; Table 1; Fig. 2D), and decrease in time constant (neonate: 125.1±30.3 ms; juvenile: 70.5±21.2 ms, *t*(_27_) = 5.956, p< 0.0001; Table 1; Fig. 2F) in RORβ interneurons from juvenile compared to neonatal cords. No significant differences were found in the rheobase (neonate: 9.1±4.4 pA; juvenile: 11.6±6.9 pA, p= 0.206; Table 1; Fig. 2C), and capacitance (neonate: 109.2±63.5 pF; juvenile: 114.3±52.7 pF, p= 0.612; Table 1; Fig. 2G). These data indicate that both the membrane potential and voltage threshold of RORβ interneurons become more hyperpolarized and input resistance decreases from neonatal to juvenile stage.

**Table 1.** Intrinsic properties of neonatal and juvenile RORβ interneurons.

| | Neonate<br>(mean±SD) | Juvenile<br>(mean±SD) | Unpaired t-test or<br>Mann-Whitney U | *p* |
| --- | --- | --- | --- | --- |
| Membrane potential (mV) | -45.1±6.4 | -50.8±6.3 | $t_{(35)}=2.754$ | 0.009* |
| Input resistance (MΩ) | 1551±911 | 718±298 | U=66 | 0.001* |
| Capacitance (pF) | 109.2±63.5 | 114.3±52.7 | U=121.5 | 0.612 |
| Time constant (ms) | 125.1±30.3 | 70.5±21.2 | $t_{(27)}=5.956$ | <0.0001* |
| Rheobase (pA) | 9.1±4.4 | 11.6±6.9 | $t_{(28)}=1.294$ | 0.206 |
| Voltage threshold (mV) | -33.1±5.5 | -38.6±4.0 | $t_{(33)}=3.526$ | 0.002* |

### Serotonin has little effect on RORβ interneurons from neonatal mice

Serotonergic projections to spinal circuits mature postnatally (31, 34, 35). To determine whether 5-HT receptors are present and activatable on RORβ interneurons at early postnatal timepoints, we first tested the effects of serotonin (5-HT) application on RORβ interneurons (*n=*9) from neonatal mice. A small but consistent depolarization was evident when 5-HT was added to the slice (baseline: −50.7±3.8 mV, 5-HT: −45.5±4.4 mV, p< 0.0001). Similarly, the resting membrane potential of RORβ interneurons depolarized in response to 5-HT (baseline: −44.0±7.7 mV, 5-HT: - 40.6±8.6 mV; p= 0.043; Table 2; Fig. 3A). However, none of the other intrinsic properties measured were significantly altered by 5-HT (Table 2; Fig. 3B-G), including rheobase (baseline: 10±4pA, 5-HT: 10.7±6.1 pA, p= 0.438), voltage threshold (baseline: −34.9±5.8 mV, 5-HT: −36.0±9.5 mV; p= 0.491), input resistance (baseline: 1199±510 MΩ, 5-HT: 1105±532 MΩ; p= 0.387), capacitance (baseline: 118±69 pF, 5-HT: 104±50 pF; p= 0.592), time constant (baseline: 116±32 ms, 5-HT: 102±50 ms; p= 0.314), and spontaneous firing (baseline: 0.17±0.24 Hz, 5-HT: 0.08±0.13 Hz; p=0.250; Fig.3 H-I) in neonatal RORβ interneurons. In RORβ interneurons which fired tonically with depolarizing current steps, there was no significant difference in the F/I slopes for neonatal RORβ interneurons (baseline: 0.36±0.05 Hz/pA, 5-HT: 0.35±0.07 Hz/pA; F=0.008; p=0.93; Fig. 3 H-I). These data show that serotonin has minimal effects on neonatal RORβ interneurons in the medial deep dorsal horn.

**Table 2.**
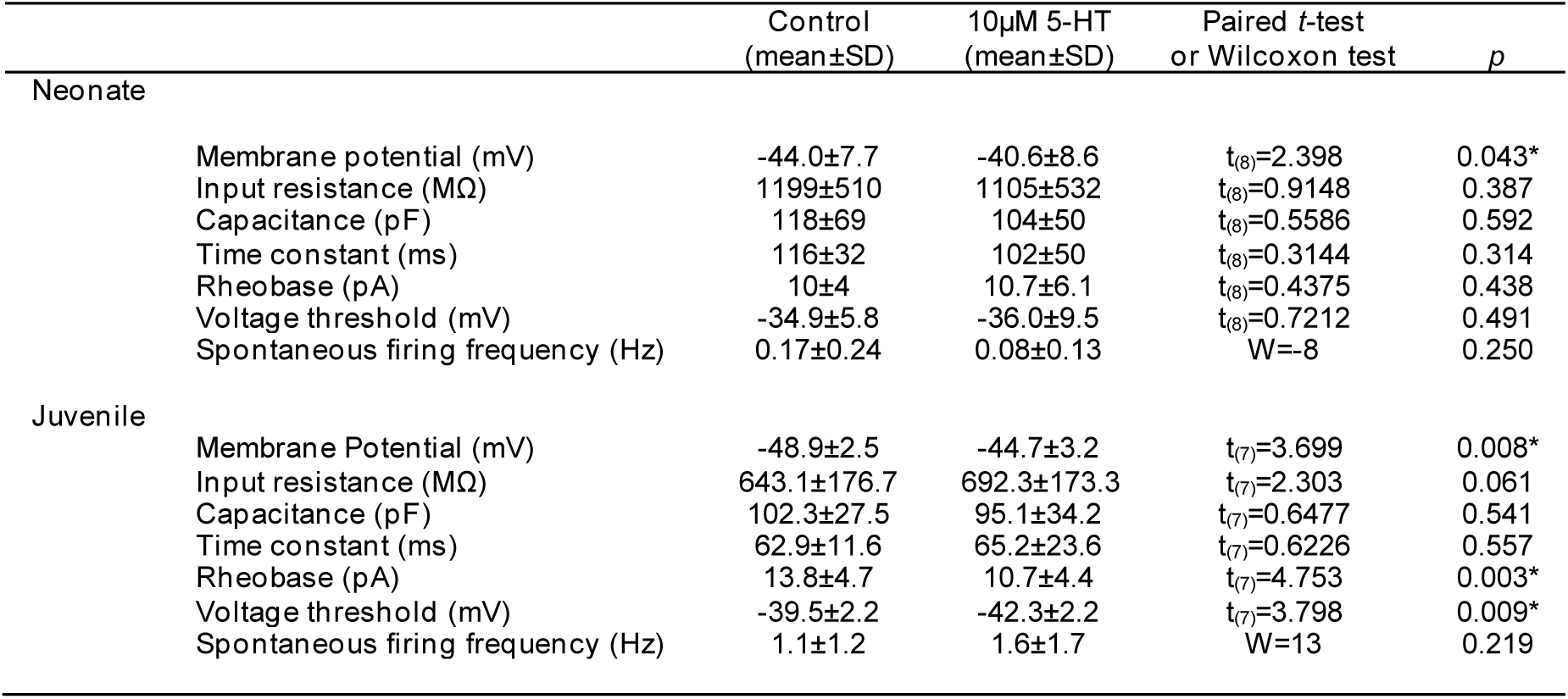
Effects of 5-HT application on neonatal and juvenile RORβ interneurons.

**Figure 3.**
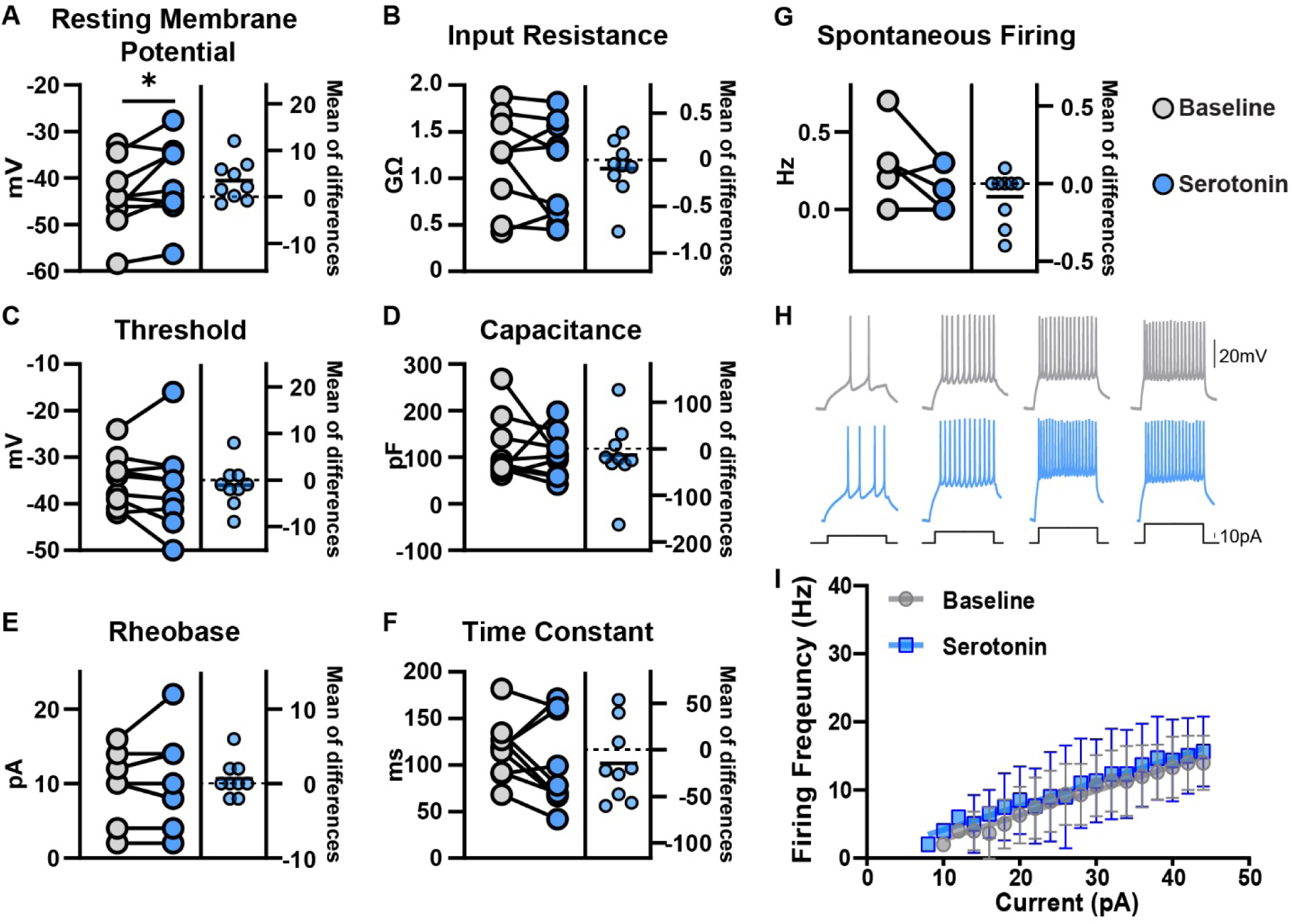
Neonatal RORβ interneurons have minimal response to serotonin. (A) Serotonin application depolarized the resting membrane potential of neonatal RORβ interneurons. (B-G) Other intrinsic electrophysiological properties did not change in response to serotonin. The rate of frequency of firing in response to depolarizing current steps (H-I) did not differ between baseline measures and in the presence of serotonin, for neonatal RORβ interneurons. Paired *t-test* or Wilcoxon matched pairs signed rank test, *=p<0.05.

### Serotonin increases the excitability of RORβ interneurons from juvenile mice

We next performed experiments to test the effects of serotonin on juvenile (*n*=8) RORβ interneurons in the medial deep dorsal horn. Addition of 5-HT to lumbar slices from juvenile mice led to an obvious depolarization of RORβ neurons (baseline: −52.8±2.3 mV, 5-HT: −47.2±3.1 mV; p< 0.0001). Similarly, resting membrane potential of RORβ interneurons was depolarized in response to 5-HT (baseline: −48.9±2.5 mV, 5-HT: −44.7±3.2 mV; p=0.008; Table 2; Fig. 4A). Although there was no significant change in input resistance (baseline: 643.1±176.7 MΩ, 5-HT: 692.3±17.33 MΩ; p= 0.061; Table 2; Fig. 4B), the rheobase was decreased by 26% when serotonin was applied to juvenile RORβ interneurons (baseline: 13.8±4.7 pA, 5-HT: 10.7±4.4 pA; p= 0.003; Table 2; Fig. 4E). Additionally, the voltage threshold for an action potential was significantly hyperpolarized in response to serotonin for juvenile RORβ interneurons (baseline: - 39.5±2.2 mV, 5-HT: −42.3±2.2 mV; p= 0.009; Table 2; Fig. 4C). Capacitance (baseline: 102.3±27.5 pF, 5-HT: 95.1±34.2 pF; p= 0.541; Table 2; Fig. 4D) and time constant (baseline: 62.9±11.6 ms, 5-HT: 65.2±23.6 ms; p= 0.557; Table 2; Fig. 4F) were not significantly altered by 5-HT. Although an increase in spontaneous firing frequency was apparent in some neurons when 5-HT was applied, it was not statistically significant for the population (baseline: 1.1±1.2 Hz, 5-HT: 1.6±1.7 Hz; p=0.219; Table 2; Fig. 4G). In juvenile RORβ interneurons which fired tonically with depolarizing current steps, there was a shift in the F/I curve with 5-HT (baseline slope: 0.52±0.04 Hz/pA, 5-HT slope: 0.67±0.04 Hz/pA; F=7.75; p=0.006; Table 2; Fig. 4H-I). These results demonstrate that the excitability of juvenile RORβ interneurons increases in response to 5-HT.

**Figure 4.**
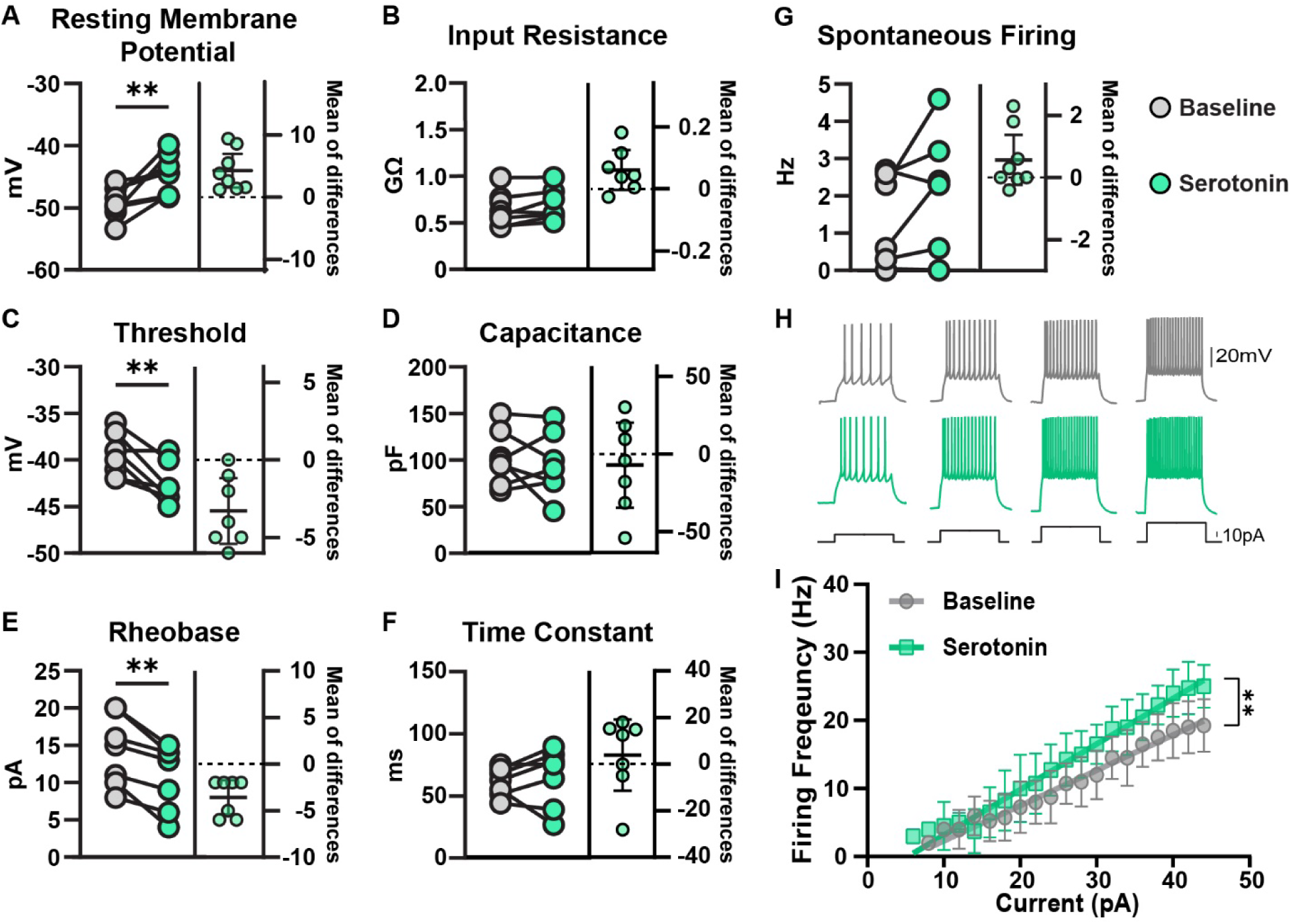
Serotonin has excitatory effects on juvenile RORβ interneurons. (A-G) Comparison of active and passive membrane properties measured at baseline (gray) and in serotonin (green) in RORβ interneurons from juvenile spinal cords. (H) Example responses to 1s long current steps in RORβ interneurons from a neonatal (gray) and a juvenile (green) mouse. (I) The slope of the frequency-current (f-I) curve was significantly steeper in the presence of serotonin. Paired t-test or Wilcoxon test,*=p<0.05, **=p<0.01; paired *t-test* or Wilcoxon matched pairs signed rank test.

### Activation of select serotonin receptors modulates the excitability of juvenile RORβ interneurons

Next, we sought to determine which serotonin receptors are functionally expressed by juvenile RORβ interneurons. Since 5-HT application had excitatory effects on juvenile RORβ interneurons, we focused on excitatory-coupled serotonin receptors expressed in the spinal cord which have been implicated in locomotion (53–56). These include 5-HT_2A_, 5-HT_2B/2C_, and 5-HT_7_ receptors, all of which have also been shown to have excitatory effects on unidentified deep dorsal horn interneurons (57). Since common 5-HT receptor agonists are not highly selective for a single receptor, we used a combination of a highly selective receptor antagonist and a receptor agonist that activates two receptor subtypes. This strategy blocked the overlapping receptor that would be co-activated by the agonist application (51). Application of selective 5-HT receptor antagonists did not affect most properties of RORβ interneurons from juvenile and neonatal cords (Supplemental Figs. 1&2 and Supplemental Tables 1&2).

We first tested the activation 5-HT_2A_ receptors on RORβ interneurons using the selective antagonist SB206553 to block 5-HT_2B/2C_ receptors (baseline) in combination with DOI, a 5-HT_2_ receptor agonist. SB206553 alone hyperpolarized the voltage threshold but had no other significant effects (Supplemental Table 1, Supplemental Fig.1). We found that activation of 5-HT_2A_ receptors on juvenile RORβ interneurons (*n*=8) caused a small (5%) but significant depolarization of resting membrane potential (baseline: −53.4±3.8 mV, 5-HT_2A_ receptor activation: −50.8±5.6 mV; p= 0.028; Fig. 5A, Table 3), as well as a 26% reduction in the rheobase (baseline: 11.3±5.3 pA, 5-HT_2A_ receptor activation 8.8±5.6 pA, p= 0.0001; Table 3; Fig. 5A). We found no significant difference in input resistance (baseline: 910±662 MΩ, 5-HT_2A_ receptor activation: 920±590 MΩ; p= 0.931; Table 3; Fig. 5A), capacitance (baseline: 108±66 pF, 5-HT_2A_ receptor activation: 105±59 pF; p= 0.824; Table 3), time constant (baseline: 68±21 ms, 5-HT_2A_ receptor activation: 74.8±27.7 ms; p= 0.239; Table 3), or voltage threshold (baseline: −37.5±5.2 mV, 5-HT_2A_ receptor activation: −38.4±5.2 mV; p= 0.088; Table 3; Fig. 5A), when activating 5-HT_2A_ receptors on juvenile RORβ interneurons. Thus, 5-HT_2A_ receptor activation both depolarizes and increases the excitability (lowers the rheobase) of RORβ neurons.

**Figure 5.**
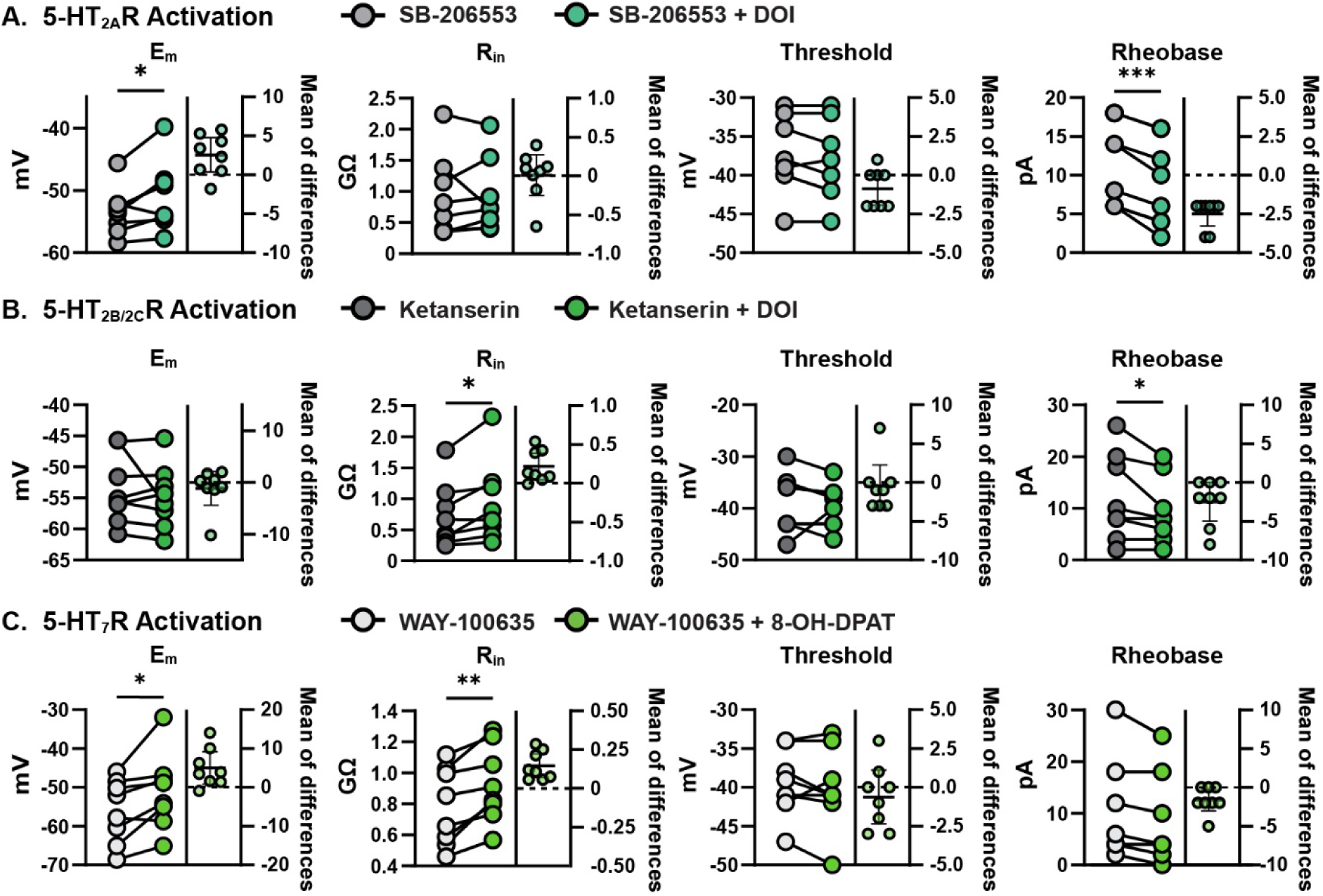
Activation of 5-HT_2A_, 5-HT_2B/2C_, and 5-HT_7_ receptors produce excitatory effects on juvenile RORβ interneurons. (A) Activation of the 5-HT_2A_ receptors on juvenile RORβ interneurons significantly depolarized resting membrane potential (E_m_) and reduced the rheobase. (B) Similarly, activation of 5-HT_2B/2C_ receptors reduced the rheobase, while increasing the input resistance (R_in_). (C) Activating 5-HT_7_ receptors on juvenile RORβ interneurons depolarized the resting membrane potential and increased the input resistance. *=p<0.05, **=p<0.01, ***=p<0.001; paired *t-test* or Wilcoxon matched pairs signed rank test.

**Table 3.** Effects of 5-HT_2A_ receptor activation on neonatal and juvenile RORβ interneurons.

|  | Neonate |  |  |  | Juvenile |  |  |  |
| --- | --- | --- | --- | --- | --- | --- | --- | --- |
| | SB206553<br>(mean±SD) | SB206553+DOI<br>(mean±SD) | Paired *t*-test<br>or Wilcoxon<br>test | *p* | SB206553<br>(mean±SD) | SB206553+DOI<br>(mean±SD) | Paired *t*-test<br>or Wilcoxon<br>test | *p* |
| *E<sub>m</sub>* (mV) | -52.3±9.7 | -52.4±9.8 | $t_{(7)}=1.874$ | 0.098 | -53.4±3.8 | -50.8±5.6 | $t_{(7)}=2.753$ | 0.028* |
| *R<sub>in</sub>* (MΩ) | 1693±669 | 1927±731 | $t_{(7)}=1.791$ | 0.116 | 910±662 | 920±590 | $t_{(7)}=0.089$ | 0.931 |
| Capacitance (pF) | 96.9±29.6 | 89.7±31.3 | $t_{(7)}=1.185$ | 0.275 | 108±65 | 105±59 | $t_{(7)}=0.231$ | 0.824 |
| Time constant (ms) | 151.6±46.2 | 151.9±48.1 | $t_{(7)}=0.065$ | 0.949 | 68±21 | 74.8±27.7 | $t_{(7)}=1.287$ | 0.239 |
| Rheobase (pA) | 9.4±4.3 | 10.2±3.7 | $t_{(7)}=0.263$ | 0.799 | 11.3±5.3 | 8.8±5.6 | $t_{(7)}=7.638$ | 0.0001* |
| Voltage threshold (mV) | -35.6±4.5 | -35.9±3.4 | $t_{(7)}=0.555$ | 0.594 | -37.5±4.9 | -38.4±5.2 | $t_{(7)}=1.986$ | 0.088 |

Next, we tested the effects of 5-HT_2B/2C_ receptor activation on juvenile RORβ interneurons *(n*=8) using the selective 5-HT_2A_ receptor antagonist ketanserin (baseline) in combination with DOI, a 5-HT_2_ receptor agonist. Ketanserin alone did not have any significant effects on RORβ interneurons (Supp. Fig. 1, Supp. Table 1). In contrast to the activation of 5-HT_2A_ receptors, the activation of 5-HT_2B/2C_ receptors on juvenile RORβ interneurons did not alter the resting membrane potential (baseline: −53.7±5.5 mV, 5-HT_2B/2C_ receptor activation: −54.8±5.1 mV; p= 0.433; Fig. 5B, Table 4). Activation of 5-HT_2B/2C_ receptors on juvenile RORβ interneurons caused a 35% increase in the input resistance (baseline: 724.9±519.7 MΩ, 5-HT_2B/2C_ receptor activation: 941.9±655.4 MΩ; p= 0.019; Table 4; Fig. 5B), and reduced the rheobase by 15% (baseline: 12±8 pA, 5-HT_2B/2C_ receptor activation: 9.5±6.3 pA; p= 0.049; Table 4; Fig. 5B). We found no significant effect on capacitance (baseline: 130.9±75.3 pF, 5-HT_2B/2C_ receptor activation: 99.7±39.8 pF; p= 0.115; Table 4), time constant (baseline: 74±32 ms, 5-HT_2B/2C_ receptor activation: 83.3±47.9 ms; p= 0.273; Table 4), and voltage threshold (baseline: −39.1±5.7 mV, 5-HT_2B/2C_ receptor activation: −39.6±4.2 mV; p=0.681; Table 4; Fig. 5B). Thus, 5-HT_2B/2C_ receptor activation also increases the excitability (decreases rheobase) of RORβ neurons, at least partly due to an increase in input resistance.

**Table 4.**
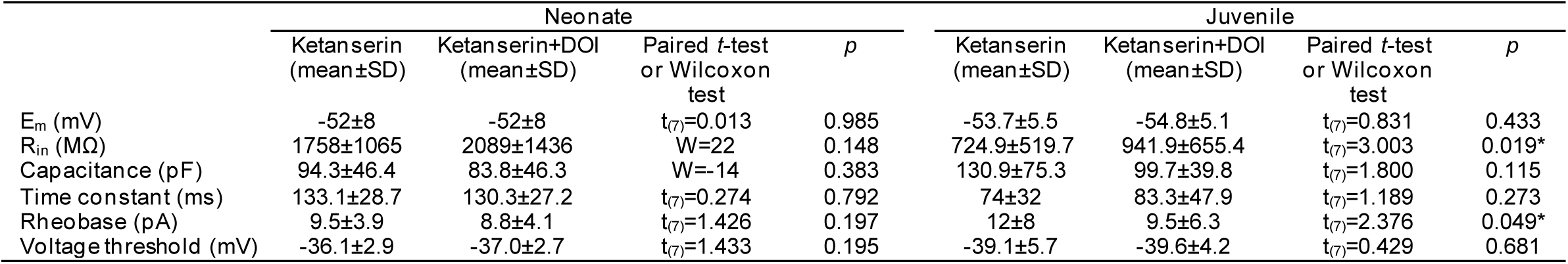
Effects of 5-HT_2B/2C_ receptor activation on neonatal and juvenile RORβ interneurons.

We then tested the activation of 5-HT_7_ receptors on juvenile RORβ interneurons by applying a selective 5-HT_1A_ receptor antagonist WAY-100635 (baseline) followed by a combination of WAY-100635 and 8-OH-DPAT which is a 5-HT_1A_ and 5-HT_7_ receptor agonist. The input resistance of RORβ neurons increased with WAY-100635 alone (Supplemental Fig. 1 and Supplemental Table 1) but all other measures were unchanged. The activation of 5-HT_7_ receptors on juvenile RORβ interneurons *(n*=8) significantly depolarized the resting membrane potential (baseline: −56.1±8.2 mV, 5-HT_7_ receptor activation: −51.1±9.8 mV; p=0.025; Table 5; Fig. 5C), and increased the input resistance by 22% (baseline: 779.1±250.2 MΩ, 5-HT_7_ receptor activation: 925±247 MΩ; p= 0.003; Table 5; Fig. 5C). There were no significant differences in capacitance (baseline: 134.6±63.5 pF, 5-HT_7_ receptor activation: 101±46 pF; p= 0.109; Table 5), time constant (baseline: 99±39 ms, 5-HT_7_ receptor activation: 89.5±33.1 ms; p= 0.437; Table 5), rheobase current (baseline: 10±10 pA, 5-HT_7_ receptor activation: 8.4±8.8 pA; p= 0.063; Table 5; Fig. 5C), or voltage threshold (baseline: −39.5±4.3 mV, 5-HT_7_ receptor activation: −40.1±5.2 mV; p= 0.421; Table 5; Fig. 5C), when activating 5-HT_7_ receptors on juvenile RORβ interneurons. Thus, 5-HT_7_ receptor activation depolarizes RORβ neurons and increases input resistance.

**Table 5.**
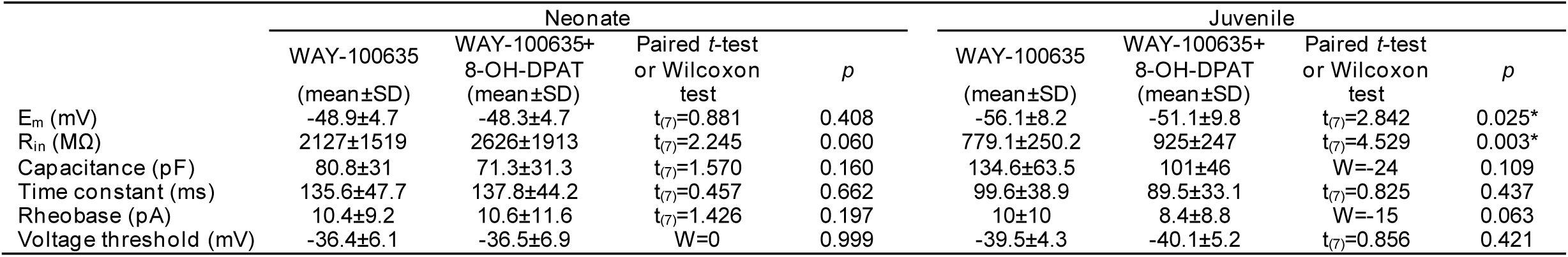
Effects of 5-HT_7_ receptor activation on neonatal and juvenile RORβ interneurons.

|  | Neonate |  |  |  | Juvenile |  |  |  |
| --- | --- | --- | --- | --- | --- | --- | --- | --- |
| | WAY-100635<br>(mean±SD) | WAY-100635+<br>8-OH-DPAT<br>(mean±SD) | Paired *t*-test<br>or Wilcoxon<br>test | *p* | WAY-100635<br>(mean±SD) | WAY-100635+<br>8-OH-DPAT<br>(mean±SD) | Paired *t*-test<br>or Wilcoxon<br>test | *p* |
| E<sub>m</sub> (mV) | -48.9±4.7 | -48.3±4.7 | t<sub>(7)</sub>=0.881 | 0.408 | -56.1±8.2 | -51.1±9.8 | t<sub>(7)</sub>=2.842 | 0.025* |
| R<sub>in</sub> (MΩ) | 2127±1519 | 2626±1913 | t<sub>(7)</sub>=2.245 | 0.060 | 779.1±250.2 | 925±247 | t<sub>(7)</sub>=4.529 | 0.003* |
| Capacitance (pF) | 80.8±31 | 71.3±31.3 | t<sub>(7)</sub>=1.570 | 0.160 | 134.6±63.5 | 101±46 | W=-24 | 0.109 |
| Time constant (ms) | 135.6±47.7 | 137.8±44.2 | t<sub>(7)</sub>=0.457 | 0.662 | 99.6±38.9 | 89.5±33.1 | t<sub>(7)</sub>=0.825 | 0.437 |
| Rheobase (pA) | 10.4±9.2 | 10.6±11.6 | t<sub>(7)</sub>=1.426 | 0.197 | 10±10 | 8.4±8.8 | W=-15 | 0.063 |
| Voltage threshold (mV) | -36.4±6.1 | -36.5±6.9 | W=0 | 0.999 | -39.5±4.3 | -40.1±5.2 | t<sub>(7)</sub>=0.856 | 0.421 |

Together, these data suggest that the 5-HT_2A_, 5-HT_2B/2C_, and 5-HT_7_ receptors are present and activatable on juvenile RORβ interneurons. Further, the excitatory effects of serotonin on juvenile RORβ interneurons are driven by a combination of 5-HT_2A_, 5-HT_2B/2C_, and 5-HT_7_ receptors.

Although 5-HT had only minor effects on RORβ neurons in neonatal slices, we tested the activation of the same receptors on neonatal RORβ neurons. None of the intrinsic properties were significantly altered by activation of 5-HT_2A_ (Table 3), 5-HT_2B/2C_ (Table 4), or 5-HT_7_ receptors (Table 5, Fig. 6). Thus, these receptors either are not present, or their second messenger cascades and/or downstream effectors are not capable of modulating the excitability of RORβ interneurons at neonatal stages. Additionally, these results demonstrate that serotonergic modulation of juvenile RORβ interneurons is mediated by serotonin 5-HT_2A_, 5-HT_2B/2C_, and 5-HT_7_ receptors and suggest that the minimal effects of serotonin on neonatal RORβ interneurons is likely due to a lack of expression of functional receptors.

**Figure 6.**
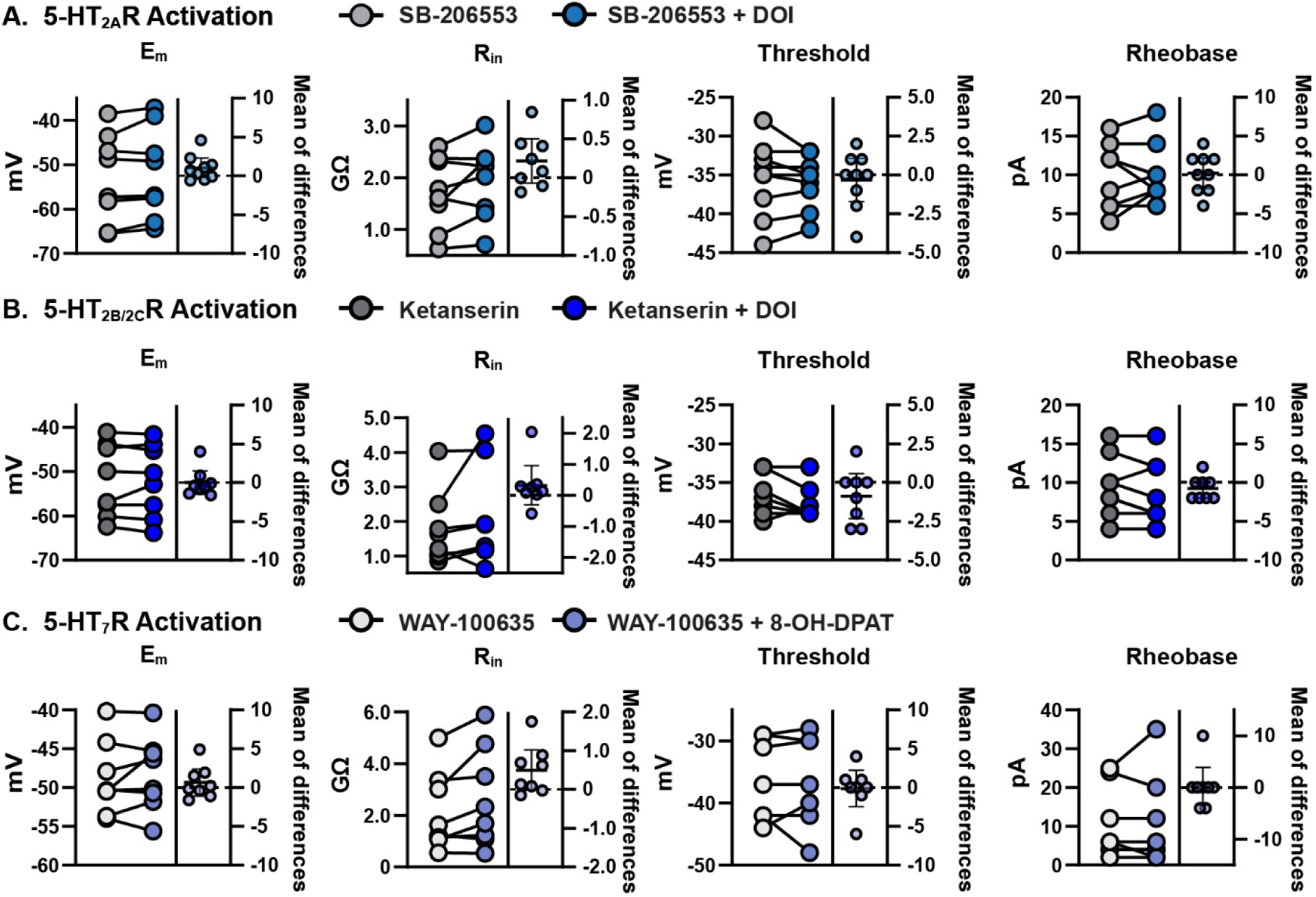
Activation of excitatory serotonin receptors has no effect on neonatal RORβ interneurons. Neonatal RORβ interneurons showed no significant change in resting membrane potential (E_m_) rheobase current, voltage threshold, or input resistance (R_in_) following activation of (A) 5-HT_2A_ receptors, (B) 5-HT_2B/2C_ receptors, or (C) 5-HT_7_ receptors Paired *t-test* or Wilcoxon matched pairs signed rank test.

## Discussion

In this study, we demonstrate that intrinsic properties of spinal deep dorsal horn RORβ interneurons mature between neonatal and juvenile stages, as does the response of these neurons to serotonin. Serotonin has excitatory actions on juvenile RORβ interneurons but only minor effects on neonatal RORβ interneurons. Further, we find that the activation of 5-HT_2_ and 5-HT_7_ receptor subtypes result in distinct excitatory effects in RORβ neurons from juvenile but not neonatal mice. Taken together, our results show that the intrinsic and neuromodulatory properties of RORβ interneurons develop postnatally and, thus, likely contribute the postnatal maturation of presynaptic inhibition.

### Deep dorsal RORβ interneurons have distinct functional and electrophysiological properties compared to superficial dorsal horn RORβ interneurons

There are two distinct populations of RORβ interneurons that can be distinguished by location in the superficial dorsal horn and medial deep dorsal horn. Medial deep dorsal RORβ interneurons mediate presynaptic inhibition of proprioceptive afferents (21), while the superficial dorsal horn population of RORβ interneurons receive mechanosensory afferent input from Aδ- and Aβ-afferent fibers (52) and are predominantly involved in postsynaptic inhibition (52). There are also electrophysiological differences between the superficial and deep dorsal horn RORβ interneurons. Delayed firing was the most prominent firing type in the superficial population (52) but delayed firing was rare in our recordings, where tonic firing was most common in both neonatal and juvenile deep dorsal RORβ interneurons. The differences in firing type may reflect a distinction of deep dorsal horn RORβ interneurons related to their role in presynaptic inhibition.

### Postnatal maturation of presynaptic inhibition is coincident with maturation of intrinsic properties of deep dorsal horn RORβ interneurons

At early developmental timepoints, inhibitory actions in the spinal cord are weak (58). Low inhibition allows for spontaneous activity to occur in spinal neurons, which is important in developing the strength of synaptic connections in spinal circuits (59, 60). However, as spinal neurons begin to mature, so do inhibitory systems, reflecting a general increase in inhibition as the spinal cord develops (60, 61). For presynaptic inhibition, part of the postnatal maturation is structural in that GAD-expressing synapses between GABAergic interneurons and proprioceptive afferents are lacking until ∼P2 and increase in number from P3-P25 (20, 28). As a result, presynaptic inhibition becomes functional postnatally after the first postnatal week (20, 28, 29).

Coincident with the establishment of presynaptic inhibition, we find that the electrical and firing properties of the deep dorsal horn RORβ interneurons mature in a similar time frame. By P14-21, a greater percentage of RORβ interneurons fire tonically compared to neonatal RORβ interneurons. The hyperpolarization of the resting membrane potential, reduction in input resistance, and decrease in time constant of juvenile RORβ interneurons compared to the neonatal RORβ interneurons are consistent with the electrophysiological changes of other spinal cord neurons occurring during the first postnatal weeks (4, 15, 50). These electrophysiological changes are occurring around the same time as the anatomical connections are maturing (20, 28, 29) and, thus, contribute to the postnatal development of functional presynaptic inhibition.

### Serotonergic modulation of RORβ interneurons develops postnatally with maturation of activatable serotonin receptors

The development of the inhibitory GABAergic phenotype in spinal neurons is influenced by serotonin (62). Serotonergic projections to the spinal cord first innervate the ventral horn as early as postnatal day 3, then later innervate the dorsal horn of the spinal cord with a notable increase at postnatal day 14 (35). Whether functional 5-HT receptor expression is established early after birth with neurons awaiting the descending projections, or whether expression follows connectivity is less clear. The latter is suggested by studies in adult spinal cord where serotonin receptor expression in the adult spinal cord is dependent on serotonergic input and/or cellular activity (63), as receptor switches and supersensitivity occur after initial receptor expression has been established (51, 64, 65). Here, we demonstrate that the serotonergic enhancement of the excitability of RORβ interneurons at juvenile ages overlaps in time with the postnatal maturation of the serotonergic projections in the area where RORβ interneurons are located (35, 66).

The neuromodulatory effects of serotonin on a given neuron are determined by the serotonin receptors expressed on the membrane of that individual neuron (57, 67, 68). We focused on 5-HT_2_ and 5-HT_7_ receptor subtypes because these receptors generally have excitatory actions (69) and they are among the primary receptor subtypes implicated in the modulation of locomotion (53, 56, 68, 70) and proprioceptive reflexes (46). 5-HT_1A_ receptors have also been implicated in locomotion and inhibition of spinal afferents (57) but these were not considered in the present study since the these receptors generally have inhibitory effects (56, 68). 5-HT_2A_, 5-HT_2B/2C,_ and 5-HT_7_ receptors have been shown to be expressed by deep dorsal horn neurons, including inhibitory neurons (57, 71), and we found evidence of functional express of all three receptor subtypes on juvenile RORβ interneurons in this study. However, we find that activating a single excitatory receptor alone did not replicate the modulatory effects of serotonin. Activation of 5-HT_2A_ and 5-HT_7_ receptors depolarized resting membrane potential. Rheobase was reduced during application of 5-HT_2A_ and 5-HT_2B/2C_ receptors, while input resistance significantly increased when 5-HT_2B/2C_ and 5-HT_7_ receptors were activated. Thus, these receptors likely contribute collectively to produce the excitatory responses seen in RORβ interneurons when serotonin is released in the spinal cord.

It is important to note that while we can record the physiological effects of activation of these receptors, there is a limitation in that we do not know the relative expression of these receptors on the membrane. It is possible that neonatal RORβ interneurons express these receptors, but they lack effect due to insufficient intracellular pathway machinery required for physiological effects (69). Additionally, it is possible that the serotonin receptors are not expressed at high enough densities on the membrane to modulate neonatal RORβ interneurons. However, considering the robust physiological effect we see when activating the excitatory serotonin receptors on juvenile RORβ interneurons, compared to the pronounced lack of effect in neonatal RORβ interneurons, we are confident that there is a meaningful difference in the expression of activatable serotonin receptors between neonatal and juvenile RORβ interneurons.

### Serotonergic modulation of RORβ interneurons plays a role in functional presynaptic inhibition

Serotonin inhibits proprioceptive afferent transmission and suppresses reflex activity through direct actions on afferents (43, 45–47), while also modulating spinal interneurons (42, 57). Previous work has speculated that serotonin reduces afferent transmission in a dual-acting mechanism, by inhibiting afferents directly (43, 47), and by increasing the activation of presynaptic inhibition (43, 72) which further suppresses afferent evoked reflexes. Given that RORβ interneurons mediate presynaptic inhibition of proprioceptive afferents (21), we directly demonstrate that serotonin increases the activity of inhibitory interneurons mediating presynaptic inhibition. The depolarization in resting membrane potential, reduced rheobase, and hyperpolarized voltage threshold seen in juvenile RORβ neurons when serotonin is present, suggests that activation of RORβ interneurons via descending serotonin contributes to the serotonergic suppression of afferent transmission (43, 56).

GABAergic presynaptic inhibition (29), descending serotonergic projections (35), and the responsiveness of RORβ interneurons to serotonin all mature by juvenile ages, around the time weight-bearing locomotion appears (4–7). Impaired development of the serotonergic system is implicated in developmental delays and sensory processing disorders (36, 37, 73). Individuals with these developmental sensory processing disorders can experience challenges with adaptation to proprioceptive demands during movement (74, 75). A lack of presynaptic inhibition contributes to sensory processing deficits associated with autism (38). Impaired development of serotonergic modulation will also impact presynaptic inhibition, with alterations in the excitability of RORβ interneurons being a contributing factor and may contribute the proprioceptive difficulties in sensory processing disorders (74, 75). Future studies may investigate changes in the activity of RORβ interneurons and serotonin receptor expression in animal models for developmental delays (36). Further, the findings we describe here can propel future investigations manipulating the serotonin receptor expression by RORβ interneurons therapeutically to improve proprioceptive difficulties in sensory processing disorders and/or following injury.

## Data Availability

All data are available in the main text or the supplementary materials.

## Acknowledgements

We thank Lihua Yao for expert technical assistance, Drexel University ULAR for animal care, and Dr. Nicholas J. Stachowski, members of the Dougherty lab, and members of the Marion Murray Spinal Cord Research Center for discussions related to the work. This work is supported by National Institutes of Health grants R01 NS104194, R01 NS130799, and T32 NS121768.

## Author contributions

C.W.W., D.L.G.-R., and K.J.D. designed research; C.W.W. performed research; C.W.W. and K.J.D. analyzed data; C.W.W. and K.J.D. wrote the paper with input from all authors.

---

**Supplementary Table 1.** Effects of serotonin receptor antagonists on juvenile RORβ interneurons. SB206553= 5-HT_2B/2C_ antagonist; Ketanserin=5-HT_2A_ antagonist; WAY-100635= 5-HT_7_ antagonist.

| | Baseline<br>(mean±SD) | SB206553<br>(mean±SD) | Paired *t*-test<br>or Wilcoxon test | *p* |
| --- | --- | --- | --- | --- |
| Membrane potential (mV) | -53.5±0.3 | -55.3±1.8 | *t*<sub>(3)</sub>=2.27 | 0.108 |
| Input resistance (MΩ) | 875.5±487.0 | 1027±447 | *t*<sub>(3)</sub>=1.29 | 0.287 |
| Rheobase (pA) | 8.5±5.3 | 6±6 | *t*<sub>(3)</sub>=1.46 | 0.239 |
| Voltage threshold (mV) | -38.8±1.3 | -40.5±1.0 | *t*<sub>(3)</sub>=1.46 | 0.035* |
| | Baseline<br>(mean±SD) | Ketanserin<br>(mean±SD) | Paired *t*-test<br>or Wilcoxon test | *p* |
| Membrane potential (mV) | -51.8±9.0 | -51.8±8.9 | *t*<sub>(2)</sub>=1.00 | 0.423 |
| Input resistance (MΩ) | 605.7±471.7 | 691±397 | *t*<sub>(2)</sub>=1.93 | 0.193 |
| Rheobase (pA) | 15.3±12.7 | 15.3±12.7 | W=-1 | 0.999 |
| Voltage threshold (mV) | -42.7±5.5 | -42±6 | *t*<sub>(2)</sub>=2.00 | 0.1835 |
| | Baseline<br>(mean±SD) | WAY-100635<br>(mean±SD) | Paired *t*-test<br>or Wilcoxon test | *p* |
| Membrane potential (mV) | -53.6±9.6 | -54.2±9.8 | *t*<sub>(5)</sub>=1.84 | 0.126 |
| Input resistance (MΩ) | 766.2±189.2 | 926.8±205.1 | *t*<sub>(5)</sub>=4.18 | 0.008* |
| Rheobase (pA) | 8.8±7.2 | 9±7 | *t*<sub>(4)</sub>=1.50 | 0.208 |
| Voltage threshold (mV) | -37.2±7.2 | -36.2±6.7 | *t*<sub>(4)</sub>=1.18 | 0.305 |

**Supplementary Table 2.**
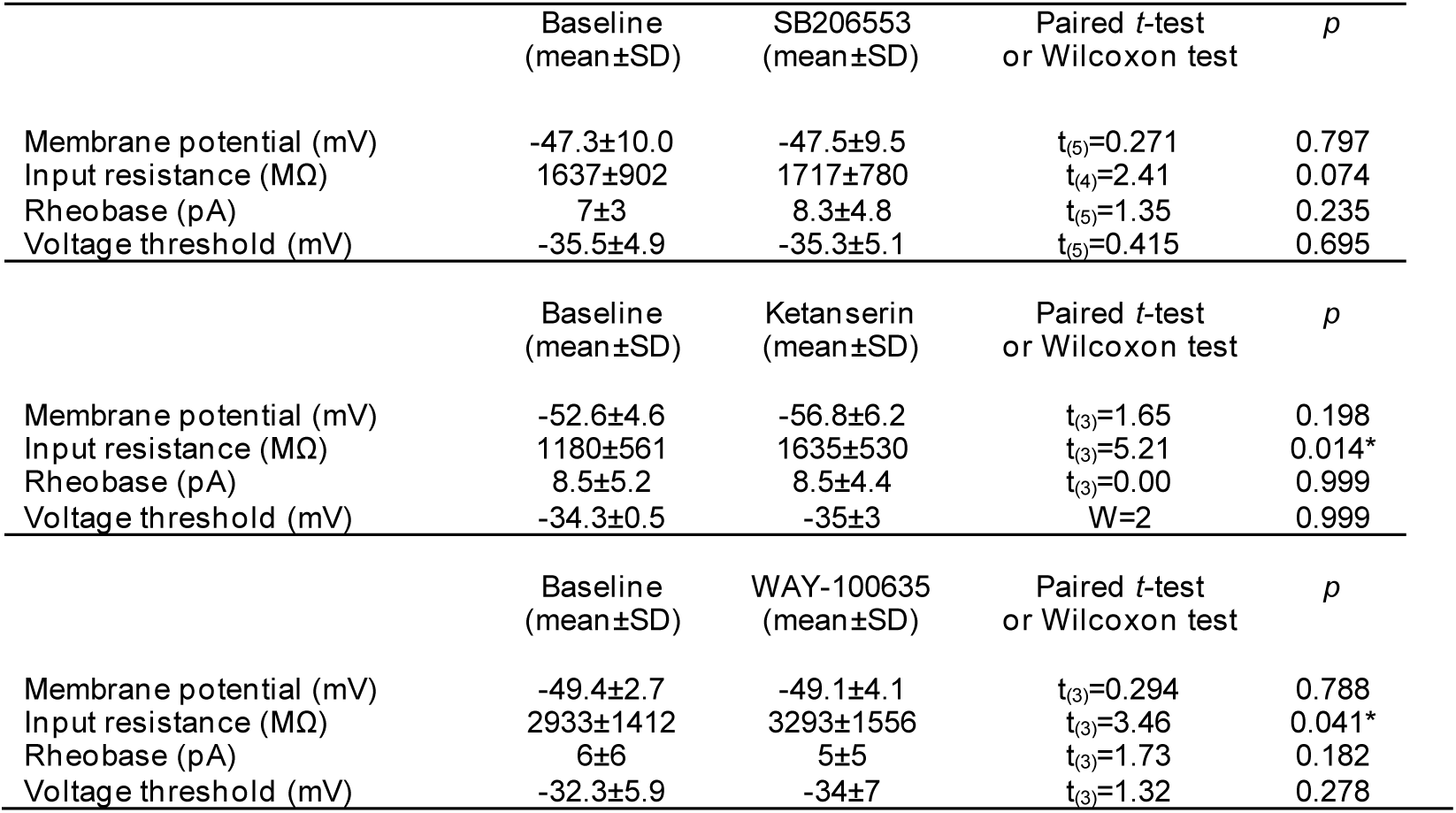
Effects of serotonin receptor antagonists on neonatal RORβ interneurons. SB206553= 5-HT_2B/2C_ antagonist; Ketanserin= 5-HT_2A_ antagonist; WAY-100635= 5-HT_7_ antagonist.

| | Baseline<br>(mean±SD) | SB206553<br>(mean±SD) | Paired *t*-test<br>or Wilcoxon test | *p* |
| --- | --- | --- | --- | --- |
| Membrane potential (mV) | -47.3±10.0 | -47.5±9.5 | *t*<sub>(5)</sub>=0.271 | 0.797 |
| Input resistance (MΩ) | 1637±902 | 1717±780 | *t*<sub>(4)</sub>=2.41 | 0.074 |
| Rheobase (pA) | 7±3 | 8.3±4.8 | *t*<sub>(5)</sub>=1.35 | 0.235 |
| Voltage threshold (mV) | -35.5±4.9 | -35.3±5.1 | *t*<sub>(5)</sub>=0.415 | 0.695 |
| | Baseline<br>(mean±SD) | Ketanserin<br>(mean±SD) | Paired *t*-test<br>or Wilcoxon test | *p* |
| Membrane potential (mV) | -52.6±4.6 | -56.8±6.2 | *t*<sub>(3)</sub>=1.65 | 0.198 |
| Input resistance (MΩ) | 1180±561 | 1635±530 | *t*<sub>(3)</sub>=5.21 | 0.014* |
| Rheobase (pA) | 8.5±5.2 | 8.5±4.4 | *t*<sub>(3)</sub>=0.00 | 0.999 |
| Voltage threshold (mV) | -34.3±0.5 | -35±3 | W=2 | 0.999 |
| | Baseline<br>(mean±SD) | WAY-100635<br>(mean±SD) | Paired *t*-test<br>or Wilcoxon test | *p* |
| Membrane potential (mV) | -49.4±2.7 | -49.1±4.1 | *t*<sub>(3)</sub>=0.294 | 0.788 |
| Input resistance (MΩ) | 2933±1412 | 3293±1556 | *t*<sub>(3)</sub>=3.46 | 0.041* |
| Rheobase (pA) | 6±6 | 5±5 | *t*<sub>(3)</sub>=1.73 | 0.182 |
| Voltage threshold (mV) | -32.3±5.9 | -34±7 | *t*<sub>(3)</sub>=1.32 | 0.278 |

**Supplementary Figure 1.**
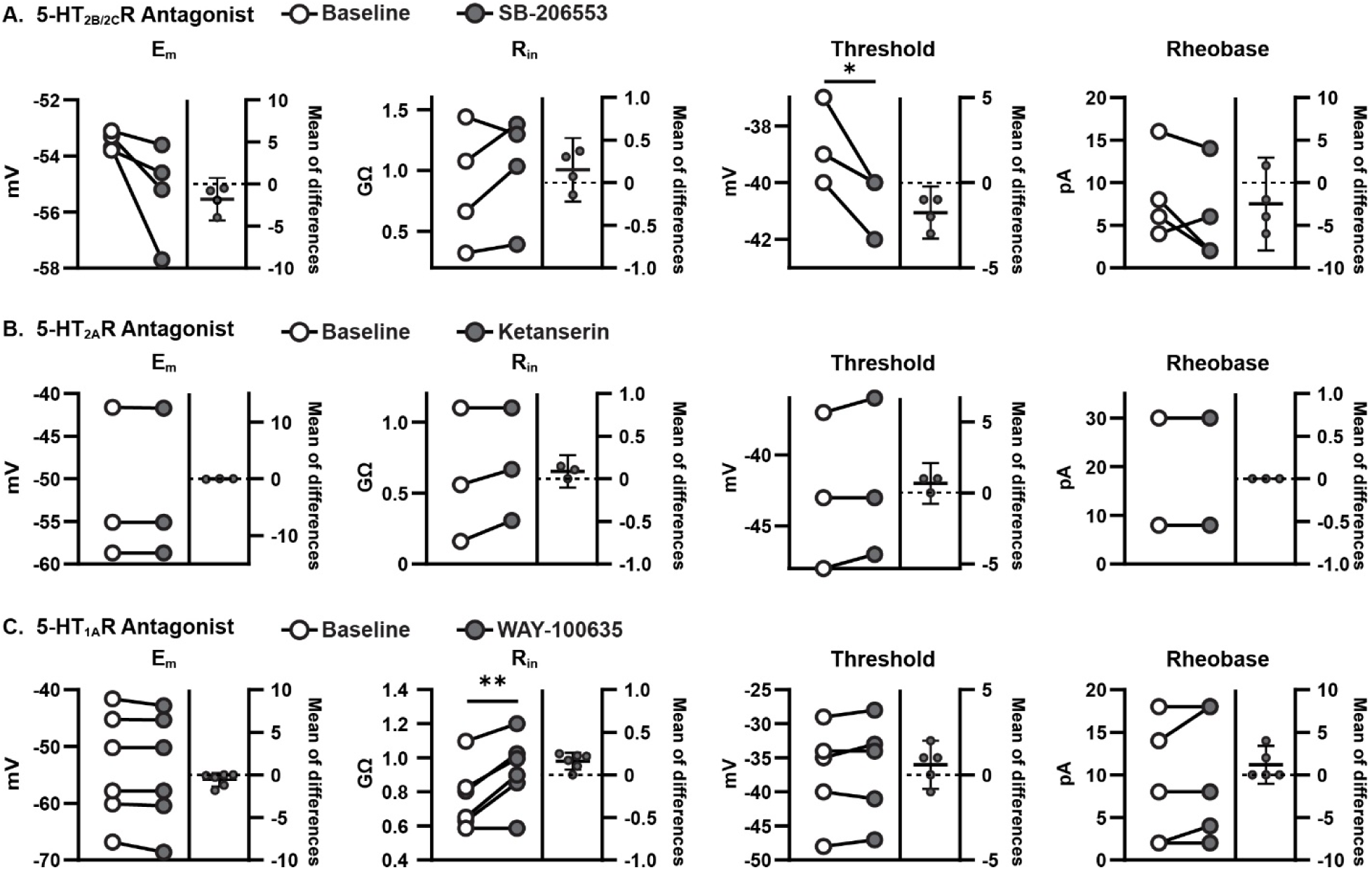
Effects of antagonists to 5-HT_2B/2C_, 5-HT_2A,_ and 5-HT_1A_ receptors on juvenile RORβ interneurons. (A) Blocking 5-HT_2B/2C_ receptors on juvenile RORβ interneurons did not significantly change resting membrane potential (E_m_), input resistance (R_in_), or rheobase. However, threshold was significantly hyperpolarized. (B) Blocking of 5-HT_2A_ receptors on juvenile RORβ interneurons also had no effect on intrinsic properties. (C) Blocking 5-HT_1A_ receptors resulted in an increase in input resistance, but no other intrinsic properties changed. *=p<0.05, **=p<0.01; paired *t-test* or Wilcoxon matched pairs signed rank test.

**Supplementary Figure 2.**
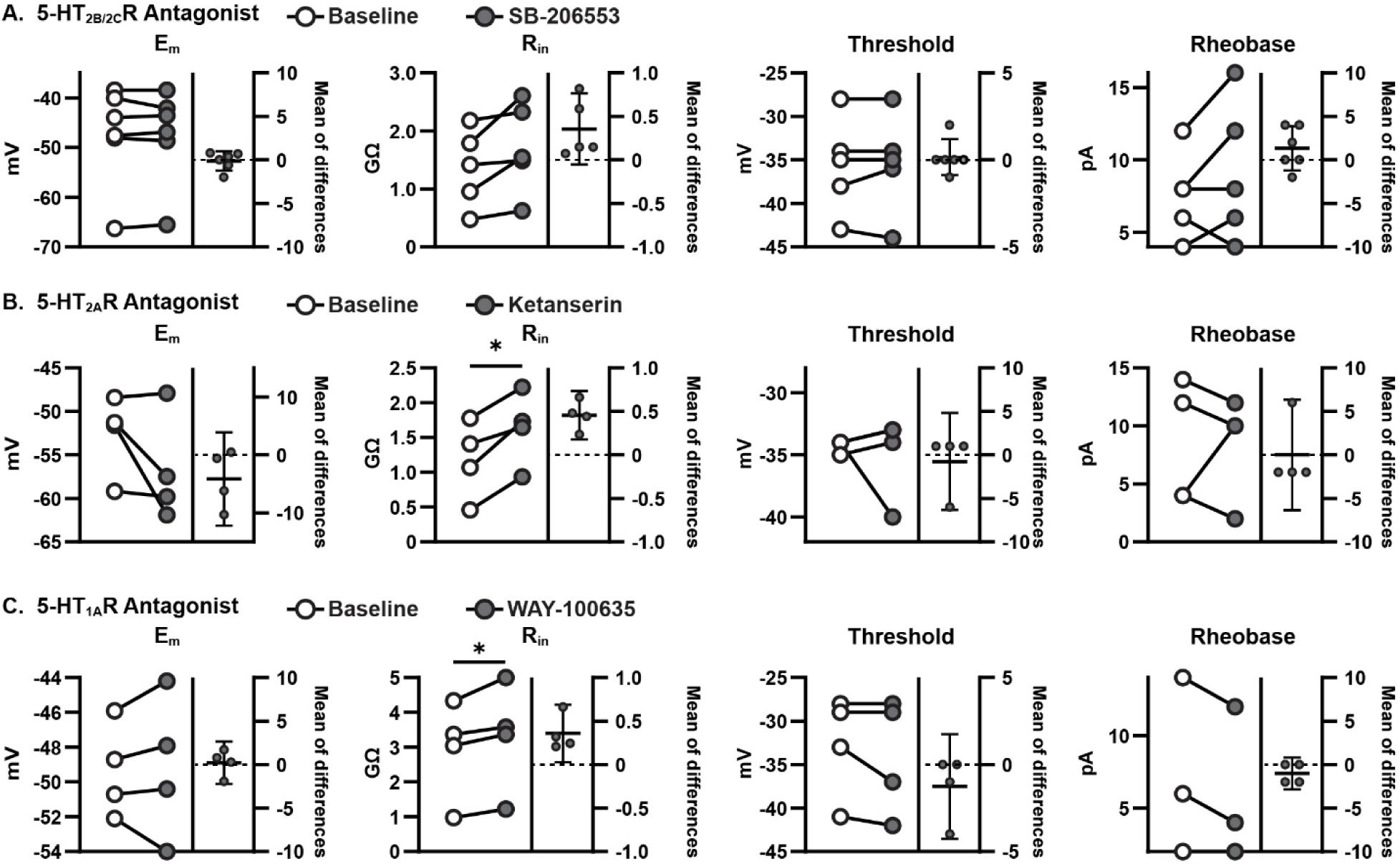
Effects of antagonists to 5-HT_2B/2C_, 5-HT_2A,_ and 5-HT_1A_ receptors on neonatal RORβ interneurons. (A) Blocking 5-HT_2B/2C_ receptors on neonatal RORβ interneurons did not significantly change resting membrane potential (E_m_), input resistance (R_in_), voltage threshold, or rheobase. (B) Blocking 5-HT_2A_ receptors significantly increased the input resistance, but no other intrinsic properties. (C) Blocking 5-HT_1A_ receptors on neonatal RORβ interneurons increased the input resistance. *=p<0.05, paired *t-test* or Wilcoxon matched pairs signed rank test.

